# *p53* deletion rescues lethal microcephaly in a mouse model with neural stem cell abscission defects

**DOI:** 10.1101/272393

**Authors:** Jessica Neville Little, Noelle D. Dwyer

**Affiliations:** Department of Cell Biology, University of Virginia School of Medicine, Charlottesville, VA, USA

## Abstract

Building a cerebral cortex of the proper size involves balancing rates and timing of neural stem cell (NSC) proliferation, neurogenesis, and cell death. The cellular mechanisms connecting genetic mutations to brain malformation phenotypes are still poorly understood. Microcephaly may result when NSC divisions are too slow, produce neurons too early, or undergo apoptosis, but the relative contributions of these cellular mechanisms to various types of microcephaly are not understood. We previously showed that mouse mutants in *Kif20b* (formerly called *Mphosph1, Mpp1*, or *KRMP1*) have small cortices that show elevated apoptosis, and defects in maturation of NSC midbodies, which mediate cytokinetic abscission. Here we test the contribution of intrinsic NSC apoptosis to brain size reduction in this lethal microcephaly model. By making double mutants with the pro-apoptotic genes *Bax* and *Trp53 (p53)*, we find that apoptosis of cortical NSCs accounts for most of the microcephaly, but that there is a significant apoptosis-independent contribution as well. Remarkably, heterozygous *p53* deletion is sufficient to fully rescue survival of the *Kif20b* mutant into adulthood. In addition, the NSC midbody maturation defects are not rescued by *p53* deletion, showing that they are either upstream of p53 activation, or in a parallel pathway. Thus, this work potentially identifies a novel midbody-mediated pathway for p53 activation, and elucidates both NSC apoptosis and abscission mechanisms that could underlie human microcephaly or other brain malformations.

## INTRODUCTION

Human genetics is increasingly successful at linking specific gene mutations to congenital brain malformations and other neurodevelopmental disorders. However, the cellular mechanisms connecting genetic mutations to brain phenotypes are still poorly understood.There are 17 human primary microcephaly genes identified, and there are many syndromes that feature microcephaly (Duerinckx and Abramowicz, 2018; Online Mendelian Inheritance in Man (OMIM) database). Thus it is a heterogeneous disorder. Known microcephaly genes encode proteins with diverse molecular functions, but many are involved in cell division.

Cell division genes may be prominent in brain malformations like microcephaly because cortical neural stem cells (NSC) divisions have several unusual features (Dwyer et al., 2016).NSCs are tall, thin cells that reside in the pseudostratified epithelium of the cortex, with their apical endfeet forming the ventricular surface, and their basal processes stretching to the basal lamina beneath the meninges. Their nuclei undergo interkinetic nuclear migration during the cell cycle, moving basally for S-phase and to the apical membrane for M-phase and cytokinesis. In addition, NSC divisions can produce symmetric or asymmetric daughter fates, giving rise to more NSCs, neurons, intermediate progenitors, and glia during corticogenesis. These stem cells must produce the right types and numbers of daughter cells within specific windows of time.With all these complex demands, the developing cortex is particularly vulnerable to insults to cell division.

We previously identified a mouse model of microcephaly that is recessive, perinatal lethal, and relatively severe, with a brain about half as thick as normal during late gestation. It carries a loss-of-function mutation in the kinesin microtubule motor gene *Kif20b*. The mutant brains do not display NSC mitotic arrest or abnormal cleavage angles, which have been noted in other microcephaly mutants. Instead, *Kif20b* mutant brains display defects in cytokinetic abscission (Dwyer et al., 2011; Janisch et al., 2013). Abscission is the process of severing the connection between mother and daughter cell, taking an hour or more after telophase (Mierzwa and Gerlich, 2014). The cleavage furrow compacts the central spindle microtubules into the midbody, which mediates abscission by recruiting proteins to remodel and “cut” the cytoskeleton and membrane. We showed that Kif20b protein localizes to the central spindle and midbody in human cell lines and mouse NSCs (Janisch, et al., 2013; 2018). Furthermore, Kif20b appears to facilitate changes in midbody microtubule structure as the midbody “matures” during the abscission process, and ensures timely abscission in cell lines. Suggesting that it may accelerate cell division, *Kif20b* is elevated in some cancers ((Kanehira et al., 2007; Liu et al.,2014; Lin et al., 2018). Interestingly, *Kif20b* evolved with the vertebrate lineage, so its subtle role in abscission may be important for growing bigger, more complex nervous systems.

In addition to abnormal midbodies, *Kif20b* mutants (*Kif20b-/-*) also display increased apoptosis in the embryonic cortex from E10.5 to E16.5 (Janisch et al., 2013). However, it was unclear whether the relatively small amount of detectable apoptosis observed could account for the severity of the microcephaly. Apoptosis appears to be relatively low in the healthy embryonic neocortex, but was seen to be elevated in some mouse models of brain malformations (Marthiens et al., 2013; Stottmann et al., 2013; Chen et al., 2014; Insolera et al., 2014; Marjanovic et al., 2015; Breuss et al., 2016). The intrinsic or stress-induced apoptotic pathway can be triggered by environmental stresses or genetic insults, such as particular mutations (Arya and White 2015). Bax and p53 (gene *Trp53*) are expressed in embryonic brain and appear to respond to damage. In response to apoptotic stimuli, Bax, a multi-BH domain-containing member of the BCL2 family, can form pores across the outer mitochondrial membrane, thereby releasing cytochrome C that can activate the caspase cascade. p53, a tumor suppressor mutated in many human cancers, is an upstream activator of Bax (Kastenhuber and Lowe 2017). Mouse knockouts of *Bax* and *p53* have normal brain development with surprisingly few low penetrance defects (Knudson et al., 1995; Jacks et al., 1994; Insolera et al., 2014).

Here, we set out to determine the relative contributions of apoptosis and abscission dysregulation to the microcephaly of the *Kif20b* mutant, and to understand the relationship between these phenotypes. To do this, we crossed genetic mutants of the intrinsic apoptosis pathway to *Kif20b*-/- mice and asked whether the apoptosis, microcephaly, and abscission defects in *Kif20b* mutants were rescued, unaffected, or potentially worsened. To our surprise, we found that *p53* deletion prevented apoptosis and rescued brain size and structure to a remarkable degree. A partial apoptosis inhibition by *Bax* deletion correlated with a lesser extent of brain size rescue. Surprisingly, deletion of even one allele of *p53* is able to completely block the apoptosis and lethality in *Kif20b* mutants. However, *Kif20b; p53* double mutant brains are still smaller than controls at birth, suggesting Kif20b regulates cortical development through additional mechanisms, perhaps by ensuring timely abscission. Indeed, *p53* deletion does not rescue the defects in midbody structure seen in *Kif20b* mutant NSCs, and additional midbody defects are revealed when *p53* is deleted. Our data provides the first evidence that abscission defects can cause p53 activation in NSCs or any cell type, through a yet-to-be identified pathway. Finally, our work addresses apoptosis inhibition as a potential way to ameliorate the severity of microcephaly caused by genetic, viral, or environmental insults.

## RESULTS

### The intrinsic apoptotic pathway is the key driver of microcephaly in *Kif20b-/-* mice

To test whether the intrinsic apoptotic pathway mediates the elevated apoptosis and microcephaly in the *Kif20b* mutant cortex, we utilized genetic crosses to mutants in two key genes in this pathway, *Bax* and *p53 (Trp53)*. Importantly, homozygous deletion mutants in *Bax* or *p53* develop properly with normal size cortices (Knudson et al., 1995; Jacks et al., 1994; Insolera et al., 2014). First, we produced *Kif20b*; *Bax* double mutants. Bax, together with its partner Bak, increases the permeability of the mitochondrial membrane, increasing cytochrome c release (Westphal et al., 2011). We examined double mutant brains at age E14.5, when apoptosis is elevated 4-fold in *Kif20b* mutants, and the cortical plate has started to form.Interestingly, embryos carrying homozygous mutations in both *Bax* and *Kif20b* showed a partial block of the elevated apoptosis and a partial rescue of cortical thickness (**Fig. 1A-H**).Craniofacial defects observed in *Kif20b* mutants were not rescued by Bax mutation (data not shown). These data suggest that apoptosis and microcephaly are correlated, but that additional proteins are required for the full apoptotic response to *Kif20b* loss. For example, Bak is a partner of Bax and is partially redundant (Lindsten et al., 2000).

**Figure 1.**
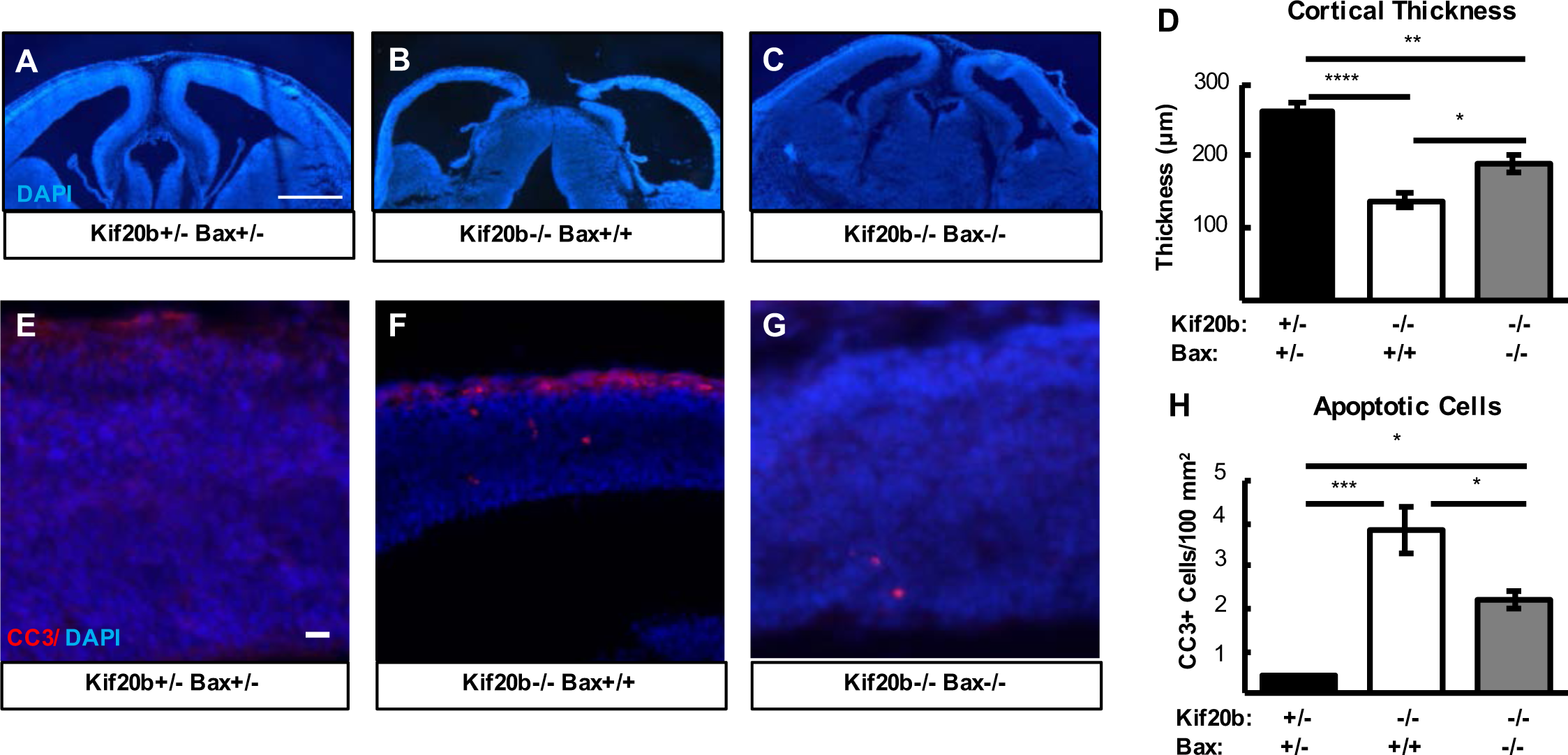
*Bax* deletion partially rescues microcephaly and apoptosis in *Kif20b-/-* mice. **A-D.** Representative E14.5 cortical sections, and plotted mean of cortical thicknesses show the severe reduction in*Kif20b* mutantsis partly restored in *Kif20b; Bax* double mutants. **E-H.** Cleaved caspase-3 (CC3) staining in E14.5 cortices shows the increased apoptosis in *Kif20b* mutants is partially reduced in *Kif20b; Bax* double mutants. For (D, H), n= 5 *Kif20b+/-; Bax+/-,* 4 *Kif20b-/-; Bax+/+*, 5 *Kif20b-/-; Bax-/-* mice, from a total of 8 litters.* p < 0.05, ** p < 0.01, *** p < 0.001,**** p < .0001, one-way ANOVA. Error bars are +/- s.e.m. Scale bars: A: 500μm; E: 20μm.

The tumor suppressor p53 is upstream of Bax/Bak in the intrinsic apoptotic pathway. Therefore, we tested whether *p53* deletion could fully block the apoptosis in *Kif20b* mutants by crossing to the *Trp53-/-* mutant (Jacks et al., 1994). Strikingly, apoptosis and cortical thickness of E14.5 *Kif20b-/-* embryos are both fully rescued by either heterozygous or homozygous deletion of *p53* (**Fig. 2A-J**). Thus, two functional *p53* genes are required to produce the apoptosis and microcephaly triggered by *Kif20b* loss. Furthermore, *p53* is required for the craniofacial defects of *Kif20b-/-* embryos: either heterozygous or homozygous *p53* deletion significantly ameliorates the small eye and snout phenotypes (**Fig. S1**). p53 protein appears to be elevated in *Kif20b* mutant cortices, as immunoblots of E12.5 cortical lysates show p53 band intensity increased 50% in *Kif20b-/-* samples, normalized to the NSC protein beta-catenin (**Fig. S2 A, B**). Furthermore, immunostaining suggests this increase is specifically in the nuclei of NSCs (**Fig S2 C, D**). Together, these data show that p53 is activated when *Kif20b* is lost, and p53 function is required for the excess apoptosis and microcephaly in this mutant. Moreover, the partial and full rescues of microcephaly by *Bax* and *p53* deletion, respectively, show that that the amount of apoptosis and the severity of microcephaly are strongly correlated, suggesting that apoptosis is the key cellular mechanism driving microcephaly in *Kif20b-/-* mice.

**Figure 2.**
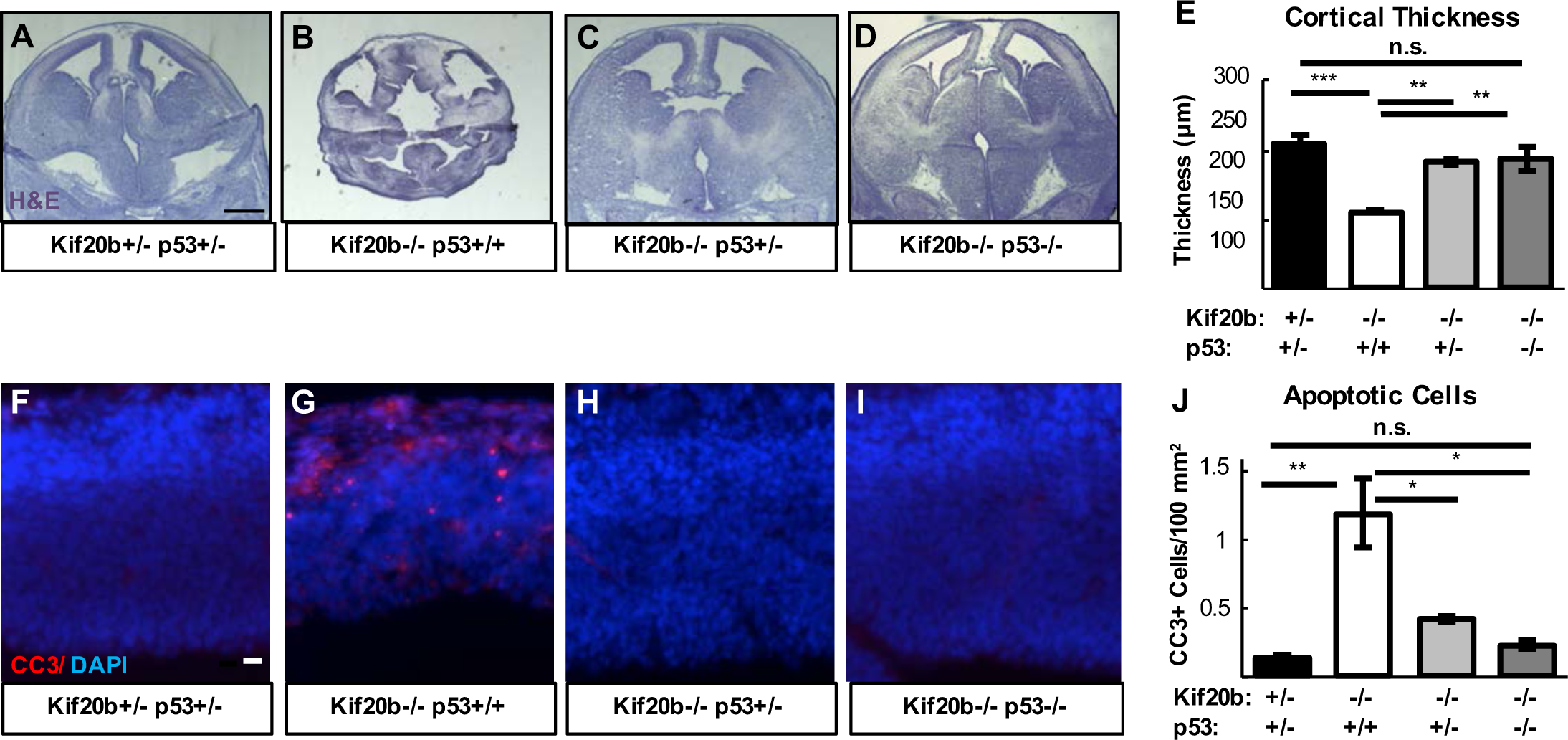
*p53* deletion rescues microcephaly and apoptosis in *Kif20b-/-* mice. **A-E.** E14.5 cortical sections and plotted mean cortical thicknesses show the severe reduction in *Kif20b* mutants is fully rescued by heterozygous or homozygous *p53* deletion. n = 6 *Kif20b+/-; p53+/-,* 3 *Kif20b-/-; p53+/+,* 4 *Kif20b-/-; p53+/-*, 4 *Kif20b-/-; p53-/-* mice, from a total of 7 litters. **F-J.** CC3 staining shows apoptosis in *Kif20b* mutants returns to control levels by heterozygous or homozygous *p53*deletion. n = 3 *Kif20b+/-; p53+/-, Kif20b-/-; p53+/+, Kif20b-/-; p53+/-, Kif20b-/-; p53-/-* mice, from a total of 6 litters. p < 0.05, ** p < 0.01,*** p < 0.001, one-way ANOVA. Error bars are +/- s.e.m. Scale bars: A: 1 mm; F: 20μm.

### *p53* deletion restores growth of neuronal and subventricular layers in embryonic *Kif20b-/-* cortex

We previously showed that the reduced cortical thickness *of Kif20b* mutants is due to thinner neuronal layers and fewer intermediate progenitors (IPs) (Janisch et al., 2013). To determine if *p53* deletion could rescue cortical neurogenesis and structure as well as thickness, we labeled *Kif20b-/-* single mutant and *Kif20b*; *p53* double mutant E14.5 cortical sections for Pax6, Tuj1, and Tbr2, to label NSCs, neurons, and IPs, respectively. In *Kif20b-/-* single mutant brains, as expected, the neuronal layer (cortical plate) and axonal layer (intermediate zone) are thin (**Fig. 3A, B, G**). Remarkably, *Kif20b-/-; p53-/-* embryos have cortices that appear to have normal organization, with cortical plates and intermediate zones indistinguishable from controls (**Fig. 3C, G**). Additionally, *Kif20b-/-; p53-/-* embryos display normal ventricular zone thickness and IP generation (**Fig. 3D-F, H, I**). Therefore, blocking apoptosis by *p53* deletion in the cortices of *Kif20b-/-* brains increased cortical thickness by improving production or survival of neurons and IPs. Furthermore, the rescued neurons and IPs appear to migrate normally and create a normal-appearing structure.

**Figure 3.**
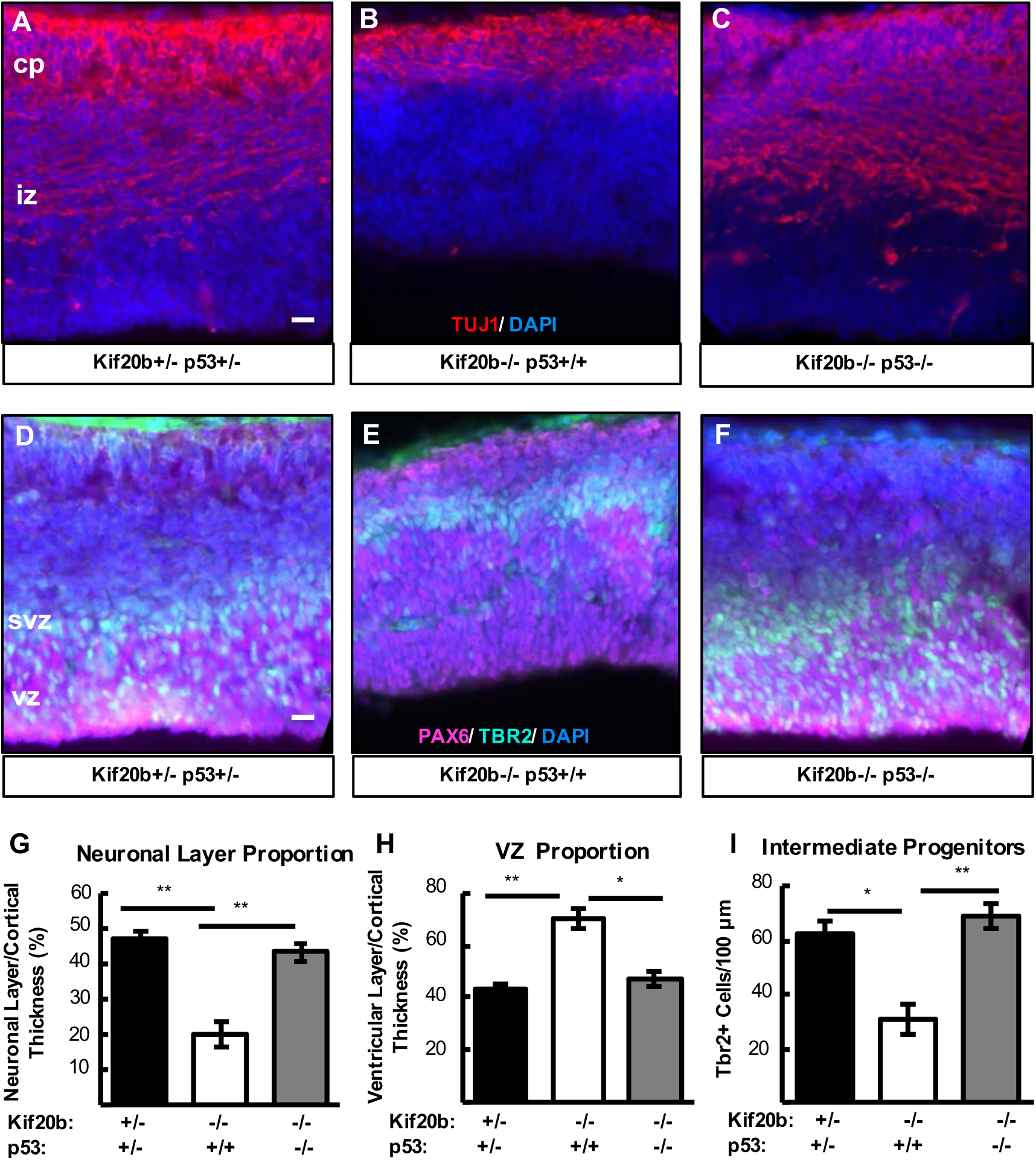
*p53* deletion restores normal production of intermediate progenitors and neurons in *Kif20b-/-* cortex. **A-C.** E14.5 cortical sections labeled with neuronal-specific tubulin (Tuj1, red) show the decreased neuronal layer thickness in *Kif20b* mutants is rescued in *Kif20b;p53* double mutants. cp: cortical plate; iz: intermediate zone. **D-F.** E14.5 cortices labeled with Pax6 (NSCs, pink) and Tbr2 (IPs, green) show the altered layer proportions in *Kif20b*mutant are rescued by *p53* deletion. svz: subventricular zone; vz: ventricular zone. **G.** The mean neuronal layer proportional thickness is halved in *Kif20b* mutantsbutrescued by *p53* co-deletion. **H.** The proper NSC layer (Pax6+) proportionality is restored by *p53* co-deletion. **I.** The density of Tbr2+ I IPs is reduced in *Kif20b* mutants, butrescued in *Kif20b;p53* double mutants. n = 3 each *Kif20b+/-;p53+/-, Kif20b-/-;p53+/+, Kif20b-/-;p53-/-* mice for Tuj1 and Pax6 layer thickness. n = 6 *Kif20b+/-;p53+/-,* 3 *Kif20b-/-;p53+/+,* 4 *Kif20b-/-;p53-/-* mice for Tbr2+ cell counts, from a total of 6 litters.p < 0.05, ** p < 0.01, one-way ANOVA. Error bars are +/- s.e.m. Scale bars: 20 μm.

### *p53* deletion rescues postnatal survival but not full cortical size of *Kif20b-/-* mice at birth

In addition to reduced brain size, *Kif20b-/-* mice exhibit perinatal lethality. Remarkably, while no *Kif20b* mutant mice with wild-type *p53* status survive the day of birth (postnatal day 0, “P0”), *Kif20b-/-* mice with heterozygous or homozygous *p53* deletion survive postnatally at expected Mendelian ratios (**Figure 4A**). Even more surprising, most *Kif20-/-; p53+/-* mice and *Kif20b-/-; p53-/-* mice live to adulthood and are fertile. Some *Kif20b-/-; p53+/-* mice (∼15%) still have visible craniofacial defects including a small eye or short snout, and ∼5% have hydrocephalus, but the majority have normal facial structure. Double homozygotes (*Kif20b-/-; p53-/-*) have even fewer craniofacial defects, but die prematurely at ∼ 3 - 4 months of age due to spontaneous tumors, as do *p53-/-* single mutants (Jacks et al., 1994).

**Figure 4.**
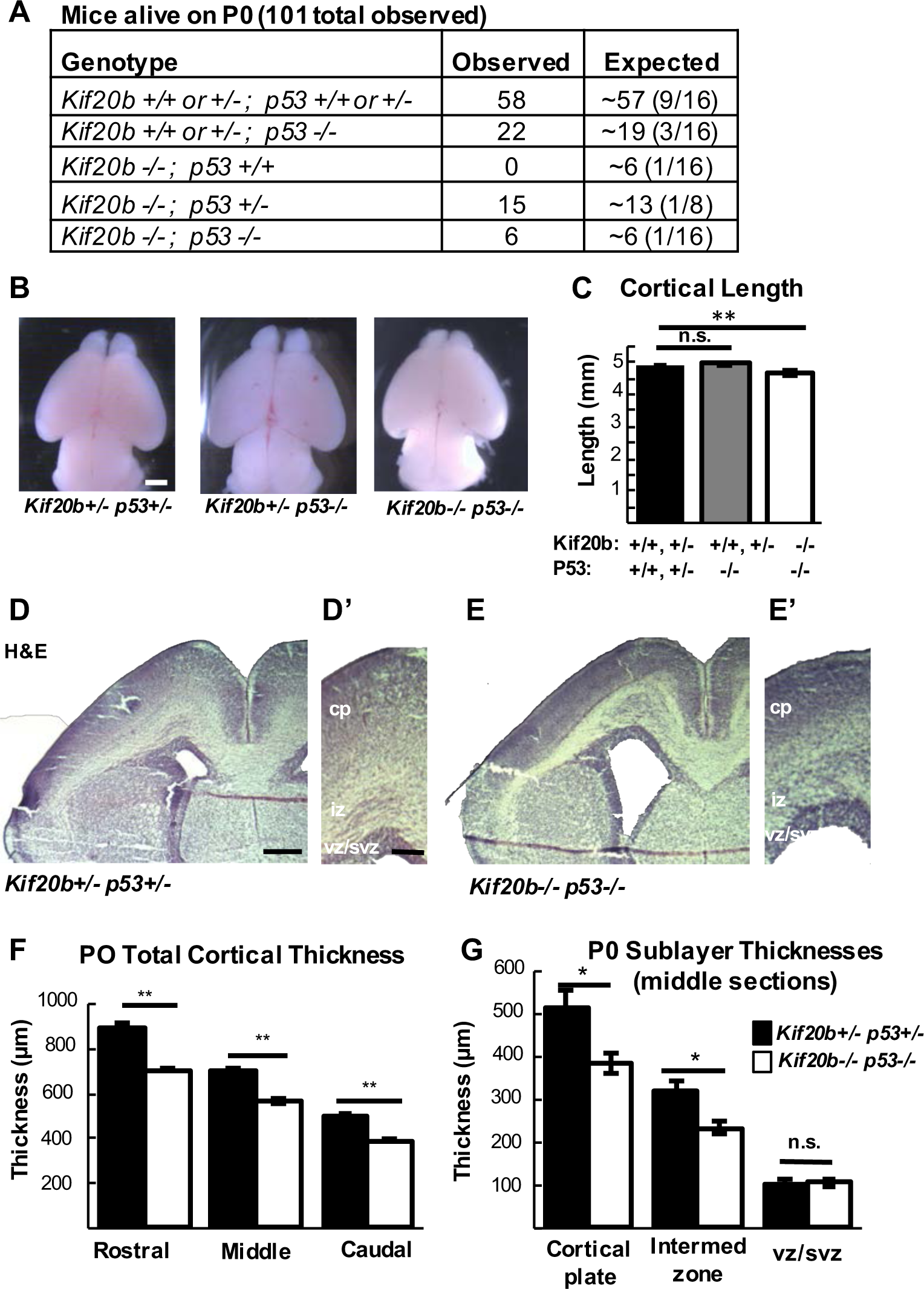
*p53* deletion rescues postnatal survival but incompletely rescues cortical size at birth in *Kif20b-/-* mice. **A.** *Kif20b-/-; p53+/+* mice do not survive atbirth at expected Mendelian ratios, but *Kif20b-/-; p53+/-* and*Kif20b-/-; p53-/-* mice survive at comparable levels to controls. **B,C.** The average length of *Kif20b-/-;p53-/-* mice is slightly decreased compared to controls at P0. N = 54*Kif20b+/+ or +/- p53+/+ or +/-* mice, 22 *Kif20b+/+ or +/- p53-/-,* and 5 *Kif20b-/-; p53-/-* mice. **D, E.** Coronal sections from P0 rostral forebrains stained with hematoxylin and eosin (H&E) show reduced cortical thickness and enlarged ventricles of double mutants. **F.** Cortical thickness in *Kif20b-/-;p53-/-* mice is reduced approximately 20% compared to controls in rostral,middle and caudal sections at P0. **G.** Reduction is in the cortical plate (cp) and intermediate zone (iz) but not ventricular zone/subventricular zone (vz/svz) layers. For F and G N = 8 *Kif20b+/-; p53+/-* and 5 *Kif20b-/-; p53-/-* brains.* p < 0.05, ** p < 0.01, one-way ANOVA (C) and student’s t-test (F,G). Error bars are +/- s.e.m. Scale bars: B:1 mm; D: 200 μm; D’: 100 μm.

The postnatal survival of *Kif20b-/-; p53-/-* mice enabled us to investigate the requirement of *Kif20b* for corticogenesis without the confounding factor of excess apoptosis. First, we confirmed previous reports that *p53* knockout mice have normal cortical size at P0 (**Fig. 4 B, C**) (Insolera et al., 2014). Interestingly, in contrast to the full rescue of cortical thickness seen at E14.5 in *Kif20b-/-; p53-/-* brains, at P0 they show a 10% decrease in cortical length and a 20% reduction in cortical thickness compared to heterozygous controls (**Fig. 4 B-F**). Furthermore, the reduction is in the cortical plate and intermediate zone, but not the ventricular and subventricular zones (**Fig. 4G**). These data indicate that despite the dramatic improvements in cortical growth and postnatal survival afforded by blocking p53-dependent apoptosis, a deficit in corticogenesis remains. Thus, while elevated apoptosis largely accounts for the microcephaly of the *Kif20b* mutant, proper cortical growth requires a *Kif20b* function that cannot be compensated by preventing apoptosis.

### *Kif20b* is required cell-autonomously for midbody maturation of cortical NSCs

Previously we showed that *Kif20b* is expressed in germinal zones of the embryonic brain, and that Kif20b protein localizes to midbodies of dividing embryonic cortical neural stem cells at the ventricular surface (Janisch et al., 2013). We further showed that *Kif20b* mutants have abnormalities in NSC midbodies: they tend to be wider and less aligned to the apical membrane. These phenotypes could be due to a cell-autonomous requirement for Kif20b during abscission, or due to non-cell autonomous effects through cell-cell interactions at the apical membrane junctions, because cytokinesis in epithelia is a multicellular process (Herszterg et al., 2014). To further probe Kif20b’s function in embryonic NSC division, we used dissociated cortical cell cultures.

Midbodies at various stages of maturation can be detected with tubulin and Aurora Kinase B (AurKB) staining by their characteristic shapes (**Fig. 5A-C**). Early-stage midbodies are wide (**Fig. 5A**), but become thinner as the midbody matures (Guizetti et al., 2011). At late stages, microtubule constriction sites (abscission sites) are detectable on one or both sides of the midbody center (**Fig. 5, B, C**, arrows) (Mierzwa and Gerlich et al., 2014). We found in human cell lines that Kif20b is recruited to early stage midbodies, and at late stages accumulates around the constriction sites (Janisch et al., 2018). Interestingly, *Kif20b-/-* NSC cultures have an increased frequency of wide midbodies (**Fig. 5D**), and fewer midbodies with at least one constriction site (**Fig. 5E**). These analyses show that *Kif20b* is required for normal NSC midbody maturation, and that this requirement is cell-autonomous.

**Figure 5.**
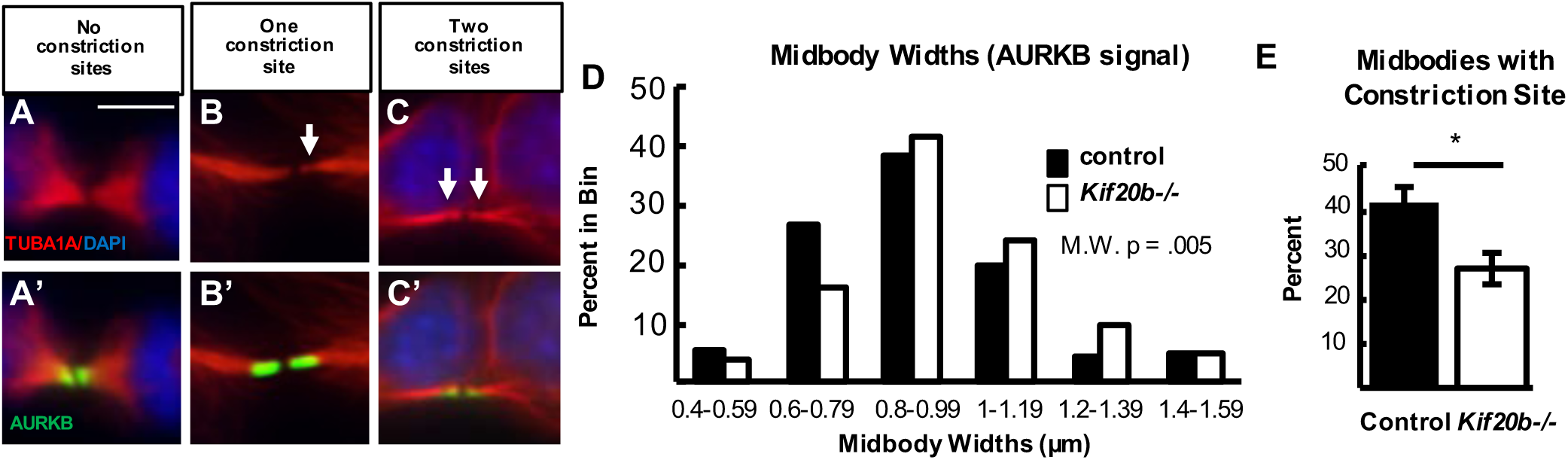
Kif20b loss cell-autonomously causes abscission defects in cortical NSCs. **A-C.** Dissociated E12.5 NSC midbodies at 24 hrs in culture labeled with alpha-tubulin (red) and Aurora kinase B (AurkB, green). Early-stage midbodies are wider with no constriction sites (A); late-stage midbodies are thinner and have one (B) or two (C) constriction sites (arrows). **D.** *Kif20b-/-* midbodies have a significantly shifted width distribution, with more wide midbodies. (Medians: Control, 0.89μm; mutant, 0.96μm). **E.** The mean percentage of midbodies with at least one constriction site (detected by tubulin) is significantly reduced in *Kif20b-/-*cultures.n (D,E): 141 control,125 mutant midbodies; 6 coverslips and 3 embryos each, 3 litters. D, Mann-Whitney; E, Student’s t-test. Error bars are +/- s.e.m. All cultures from E12.5 cortices and fixed after 24 hours. Scale bar in A: 5 μm

Some other mouse models of microcephaly show increased mitotic index in the cortex, due to NSC mitosis delay or arrest (Marthiens et al., 2013; Chen et al., 2014; Insolera et al., 2014; Marjanovic et al., 2015; Breuss et al., 2016). We showed previously that the *Kif20b* mutant cortex does not have increased mitotic index (Janisch et al., 2013), but here we use the dissociated cortical cell cultures to address this possibility with higher cellular resolution. We assayed whether *Kif20b* mutant NSCs spent more time in mitosis or abscission than control NSCs by determining the mitotic and midbody index in cultures. Surprisingly, among cycling NSCs (Ki67+) in *Kif20b* mutant cultures, both the mitotic index and midbody index were not increased but were actually slightly reduced (**Fig. S3B**). This could suggest that *Kif20b* mutant NSCs undergo mitosis and abscission more rapidly than control cells, or that some undergo apoptosis. To analyze relative durations of cell division phases, we categorized mitotic and midbody stage NSCs into sub-stages by PH3 and AurKB appearance (**Fig. S3A, C**). Among mutant NSCs at some stage of cell division, the percentages in prophase or prometa/metaphase were not different, but the percentage in anaphase/early telophase was slightly increased in *Kif20b* mutant cultures. These data are consistent with our previous results that early steps of cell division are not disrupted in *Kif20b* mutant NSCs, but that cytokinesis is affected (Janisch et al 2018).

### *Kif20b* loss activates apoptosis cell-autonomously in proliferating cortical NSCs

An attractive hypothesis is that defective midbody maturation of NSCs in *Kif20b* mutants activates the intrinsic pathway of apoptosis in NSCs. However, our previous *in vivo* analysis could not distinguish whether the apoptotic cells in *Kif20b* mutant brains were NSCs or neurons, and whether apoptosis was triggered autonomously or by cell-cell interactions. Therefore, we used the dissociated cultures to distinguish these possibilities. Indeed, in cultures of *Kif20b* mutant cells after one day *in vitro*, apoptosis is elevated more than two-fold over controls (**Fig. 6A, B, E;** arrows, CC3+ cells), suggesting that apoptosis is triggered autonomously.

**Figure 6.**
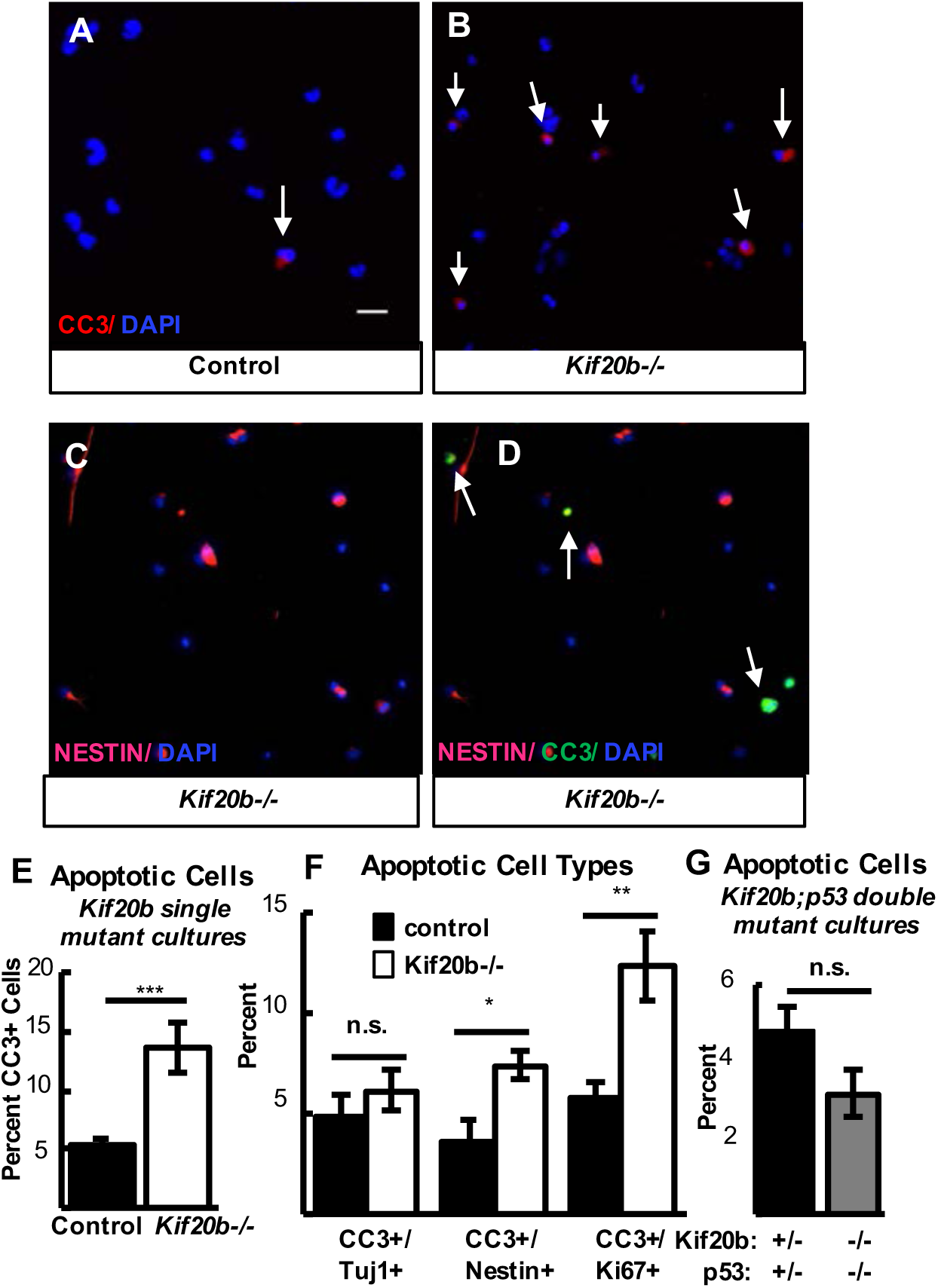
*Kif20b* loss causes apoptosis in proliferating cortical NSCs. **A,B.** Representative fields of control (left) and *Kif20b-/-* (right) dissociated cortical cultures labeled with CC3 (red, arrows) for apoptotic cells. **C,D.** Representative fields labeled with Nestin (NSCs, pink) and CC3 (green). Arrows in D indicate cells co-positive for Nestin/CC3. **E.** The mean percentage of apoptotic cells in *Kif20b-/-* cortical cultures is increased (> 2 fold) above controls. n = 13 control, 11 mutant coverslips; 8 and 7 brains, 7 litters. **F.** The percent of apoptotic neurons (CC3+/Tuj1) is not increased in *Kif20b-/-* cultures, butthe percents of apoptotic NSCs and proliferating NSCs (CC3+/Nestin+; CC3+/Ki67+) are significantly increased. Tuj1/CC3 analysis: n = 7 control, 6 mutant coverslips; 4 and 3 brains, 4 litters. Nestin/CC3 analysis: n = 7 control, mutant coverslips; 4 brains each, 3 litters. Ki67/CC3 analysis: n = 6 control, 5 mutant coverslips; 3 brains each, 3 litters. **G.** Apoptosis is not increased in *Kif20b;p53* double mutant cultures. n = 4 *Kif20b+/-;p53+/-*, 3 *Kif20b-/-;p53-/-* coverslips; 3 and 2 embryos, 2 litters. * p < 0.05,** p < 0.01,*** p < 0.001, Student’s t-test. Error bars are +/- s.e.m. All cultures from E12.5 cortices and fixed after 24 hours. Scale bars in A: 20 μm.

Furthermore, neurons (TuJ1+) do not display increased apoptosis, while NSCs (Nestin+) do, and cycling NSCs (Ki67+) show a more pronounced increase (**Fig. 6 C, D, F**). This may explain the reduced mitotic and midbody indices among mutant NSCs (**Fig. S2B**), as some arrest in the cell cycle or die. Finally, since the elevated apoptosis in *Kif20b-/-* brains requires *p53* (**Fig. 2**), we wondered whether dissociated *Kif20b-/-* NSC apoptosis also requires p53. Indeed, in cultures from double mutant mice, the rate of apoptosis is similar to controls, showing that the elevated apoptosis in isolated *Kif20b-/-* NSCs is also p53-dependent **(Figure 6G).** Together, these data show that loss of *Kif20b* causes cell-intrinsic, p53-dependent apoptosis, specifically in cycling NSCs but not postmitotic neurons.

### *p53* deletion does not rescue impaired abscission in *Kif20b* mutant mice

The preceding data show that in *Kif20b* mutant brains, the NSC apoptosis and microcephaly are downstream of p53 activation. However, another important question is whether the midbody defects seen in Kif20b mutant mice, indicating an abnormal abscission process, are upstream or downstream of p53 activation. Defects or delays in midbody maturation could activate p53. Alternatively, these midbody defects could be a consequence of p53 activation, since p53 can regulate many genes and processes. To distinguish these possibilities, we tested whether abnormal midbody phenotypes observed in *Kif20b* mutant NSCs are rescued by *p53* deletion. First, we analyzed NSC midbody index in the dissociated cultures. As *Kif20b-/-* NSCs *in vitro* are less frequently observed at the midbody stage **(Figure S3B),** and we hypothesized this is due to NSC arrest or apoptosis, we predicted this phenotype would be rescued by *p53* deletion. In fact, the midbody index of *Kif20b-/-; p53-/-* NSCs is not merely rescued, but instead is significantly increased above controls **(Fig. 7A, B).** This is consistent with the notion that some *Kif20b-/-* midbody-stage NSCs die, and further suggests that if these cells are prevented from dying, they take longer to complete abscission than control cells. Next, we analyzed NSC midbody structure in dissociated cultures. Similar to *Kif20b-/-* midbodies, *Kif20b-/-; p53-/-* midbodies are significantly wider than controls (data not shown), and fewer of them have constriction sites **(Fig. 7C).** Thus, unlike apoptosis, the midbody width and constriction site phenotypes are not p53-dependent.

**Figure 7.**
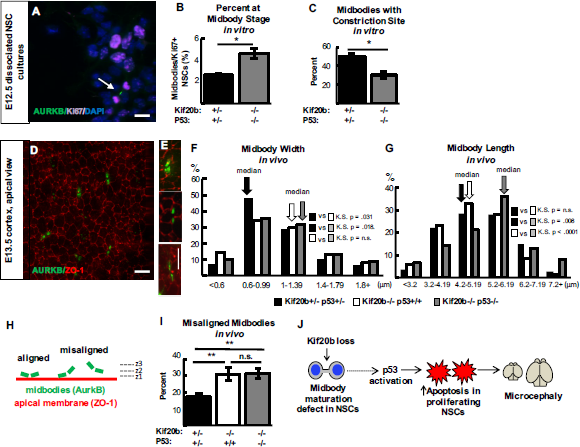
*p53* deletion does not rescue abscission phenotypes of *Kif20b* mutant NSCs. **A.** Representative image of E12.5 dissociated cortical cells, labeled with AurkB for midbodies (green, arrow), and Ki67 for proliferating NSCs (pink). **B.** The mean percent midbody stage proliferating NSCs is significantly increased in *Kif20b-/-;p53-/-* cultures. n = 4*Kif20b+/-;p53+/-*, 3 *Kif20b-/-;p53-/-* coverslips; 3 embryos each, 2 litters. **C.** Of the NSCs at midbody stage, the percent with a constriction site is significantly reduced in *Kif20b;p53* cultures. n = 111 *Kif20b+/-;p53+/-*, 101 *Kif20b-/-;p53-/-* midbodies; 3 embryos each, 2 litters. **D.** Representative image of E13.5 cortical slab, apical surface, labeled with ZO-1 (red, apical junctions) and AurkB (green, midbodies). **E.** NSC midbodies range from short and wide (top) to long and thin (bottom). **F.** The median midbody width is similarly increased in *Kif20b* single and *Kif20b;p53* double mutant cortices compared to controls. Median widths (μm) = 0.92 in *Kif20b+/-;p53+/-,* 1.0 in *Kif20b-/-;p53+/+,* 1.02 in *Kif20b-/-;p53-/-* cortices. **G.** The median midbody length is similar in *Kif20b*-/- and control cortices, but is significantly increased in *Kif20b-/-;p53-/-* cortices. Median lengths (μm) = 4.82 in *Kif20b-/-;p53+/+,* 5.11 in *Kif20b+/-;p53+/-*, 5.43 in *Kif20b-/-;p53-/-* cortices. n for (F,G) = 346 *Kif20b+/-;p53+/-,* 189 *Kif20b-/-;p53+/+*, and 299 *Kif20b-/-;p53-/-* midbodies; 8, 5, and 7 corticalhemisphere slabs, 6,4, and 5 embryos, 8 litters. **H.** Schematic ofaligned and misaligned midbodies relative to the apical membrane. **I.** The percent misaligned midbodies is doubled in both *Kif20b-/-* and *Kif20b-/-;p53-/-* cortices compared to controls. n = 8 *Kif20b+/-;p53+/-,* 7 *Kif20b-/-;p53+/+*, 7 *Kif20b-/-;p53-/-* cortical slabs from 6,4, and 6 embryos, respectively, 8 litters. **J.** Working model. p53 is required for NSC apoptosis and microcephaly caused by *Kif20b loss* but is notrequired for the midbody defects. How midbodydefects lead to p53 activation remains unknown (dotted arrow).A-C are E12.5 cortical cultures fixed at 24 hrs. *p < 0.05, ** p < 0.01, Student’s t-test for (B,C); K.S. test for (F,G). Error bars are +/- s.e.m. Scale bars, 10 μ m.

We next analyzed *Kif20b-/-; p53-/-* double mutant midbodies *in vivo* in E13.5 cortices. Cytokinesis is more complex in the cortical neuroepithelium than *in vitro.* NSC nuclei must migrate to the apical membrane to undergo mitosis and cytokinesis. During cytokinesis, the cleavage furrow ingresses asymmetrically from the basal side of the cell, forming the midbody at the apical membrane (Kosodo et al., 2008). To visualize NSC midbodies *en face*, we immunostained cortical slab whole-mounts for AurKB and the apical junction marker zona occludens-1 (ZO-1), and imaged the apical/ventricular surfaces **(Fig. 7D).** As *in vitro*, some midbodies are short and wide, and others are long and thin, since midbodies narrow as they mature (examples, **Fig. 7E**). In *Kif20b-/-* cortices, midbodies have a shifted width distribution, with an increased median width compared to controls (**Fig. 7F** and Janisch and Dwyer, 2016).

Interestingly, we find that the *Kif20b-/-; p53-/-* double mutant cortices have a strikingly similar midbody width distribution as *Kif20b* single mutants, with the same median **(Fig. 7F).** Thus, this midbody maturation phenotype (width) i*n vivo* is not rescued by *p53* deletion. But surprisingly, we find that midbody lengths, which are similar in controls and *Kif20b-/-* single mutants, are significantly longer in *Kif20b-/-; p53-/-* double mutants than in either *Kif20b-/-* single mutants or controls **(Fig. 7G)**. Exceptionally long midbodies were observed after delayed abscission in cell lines that are resistant to apoptosis (Gromley et al., 2003; Weiderhold et al., 2016). Thus, *Kif20b* mutant NSCs may have delayed abscission, which can manifest as longer midbodies only if apoptosis is prevented. An additional midbody phenotype we observe *in vivo* in *Kif20b* mutant cortices is that about a third of NSC midbodies are not aligned parallel to the apical membrane, compared to ∼15% in control brains (Janisch et al., 2013; **Fig. 7H, I**). The cause of this phenotype is unclear, but we used the *Kif20b; p53* double mutants to test whether it is p53-dependent, perhaps due to the apoptotic process in dividing or neighboring cells. Interestingly, *p53* deletion did not prevent the misalignment phenotype, showing that it is not due to p53 activation or apoptosis **(Fig 7I)**. Taken together, these midbody analyses support the hypothesis that defective midbody maturation and alignment are primary consequences of *Kif20b* loss rather than secondary consequences of p53 activation. Further, the additional midbody phenotypes detected in *Kif20b-/-; p53-/-* double mutant NSCs, namely increased midbody index and midbody length, are consistent with delayed midbody maturation in *Kif20b* mutant NSCs; however, these symptoms of delay are precluded in *Kif20b* mutants by p53 activation and apoptosis.

## DISCUSSION

Here we have tested the contribution of the intrinsic apoptosis pathway to the reduced cortex size in a lethal microcephaly model, the *Kif20b* mouse mutant. We have shown that apoptosis of cortical NSCs accounts for most of the microcephaly, but that there is a significant apoptosis-independent contribution as well, likely reflecting the importance of Kif20b’s role in abscission for corticogenesis. Furthermore, we showed that the excess apoptosis is partially dependent on Bax, and fully dependent on p53. Remarkably, heterozygous *p53* deletion is sufficient to fully suppress the lethality of the *Kif20b* mutant, and rescues the brain size equally as well as homozygous *p53* deletion. Lastly, we demonstrated that the NSC midbody maturation defects are not rescued by *p53* deletion, which indicates that they are not caused by p53 activation, but may be upstream of p53. Thus, this work potentially identifies a novel midbody-initiated pathway for p53 activation, and suggests that at least some types of microcephaly, although severe, could be greatly ameliorated by inhibiting apoptosis.

The genetic and cellular experiments herein support the following working model for the etiology of microcephaly in the *Kif20b* mutant **(Fig. 7J)**. Midbody maturation defects in some *Kif20b-/-* NSCs cause p53 activation by an unknown molecular pathway (dashed line). p53 then triggers the apoptotic cascade including Bax and other effectors. Apoptosis depletes the NSC pool, thereby reducing the number of neuron and IP daughters produced, resulting in a small brain. There is likely some stochasticity to this process, as not all *Kif20b-/-* NSCs have midbody defects, and only a small percentage undergo apoptosis (Janisch et al., 2013; this work). It may be that apoptosis is only triggered if midbody maturation (and hence abscission) is delayed beyond a certain threshold. This would be analogous to a previous study in developing cortex showing that apoptosis likelihood increases if prophase is delayed past a threshold (Pilaz et al., 2016). Though both of these types of cell division delays trigger apoptosis through p53, the molecular pathway upstream of p53 is likely distinct, since midbody maturation takes place well after telophase, in the next G1 phase (Gershony et al., 2014). In fact, how defects in early steps of mitosis signal to trigger p53 activation is only beginning to be elucidated, primarily in immortalized cell lines that are apoptosis-resistant (McKinley and Cheeseman 2017; Lambrus et al., 2016; Meitinger et al., 2016). The mechanism by which abscission delay could activate p53 or apoptosis has not been addressed in any system. Much more work is needed to determine whether and how a midbody “error sensor” could directly or indirectly activate p53 in NSCs and other cell types. It remains possible that *Kif20b* loss causes p53 activation and midbody defects through two separate pathways. In either case, this mutant is a tool to elucidate a novel pathway to p53 activation, one that appears very sensitive in cortical neural stem cells.

### p53 activates apoptosis in NSCs following diverse cellular defects

Our work furthers the evidence that p53 responds to multiple intrinsic cellular defects to acutely regulate NSC survival. Zika virus infection activates p53-dependent apoptosis in human NSCs (Ghouzzi et al., 2016). A handful of other microcephalic mouse mutants, with impaired DNA replication, mitosis, or cleavage furrowing, have implicated p53 in apoptosis and reduced brain size (Bianchi et al., 2017; Houlihan et al., 2014; Insolera et al., 2014; Marjanovic et al., 2015; Marthiens et al., 2013; Murga et al., 2009). Interestingly, inhibition of p53 in these mutants was sometimes able to increase cortical thickness, but in other cases worsened brain phenotypes (Marthiens et al., 2013; Murga et al., 2009). By contrast, *Kif20b-/-* embryonic brain structure and postnatal survival were well-rescued by even heterozygous deletion of *p53*. Heterozygous *p53* rescue has not been reported in other microcephaly mouse mutants. In the *Brca1* mutant, dwarfism but not microcephaly was rescued by heterozygous *p53* deletion (Xu et al., 2001). Thus, the *Kif20b-/-* microcephaly model is more severe in reduction of brain size compared to some other models, but it is also more easily rescuable by apoptosis inhibition. In fact, *p53* heterozygous or homozygous deletion can completely rescue the perinatal lethality of *Kif20b-/-* mice. This is important for eventual treatment of human microcephaly caused by genetic mutations or viruses, to determine which subtypes and phenotypes might be treatable by blocking apoptosis.

### A p53-independent consequence of Kif20b loss accounts for the remaining cortical size deficit at birth

The 20% deficit in cortical thickness in *Kif20b*; *p53* double mutant mice at birth indicates that even without the excess apoptosis, *Kif20b* mutant NSCs and neurons cannot create a brain of normal size and structure, probably due to abnormal abscission causing some other problem in *Kif20b-/-; p53-/-* NSCs. The midbody maturation defects that are more frequent in *Kif20b* mutant brains may cause delays in daughter cell severing from the mother cell and delamination from the apical membrane in the case of neuronal daughters. It could also affect whether abscission occurs on both sides of the midbody or only one. The significance of the midbody misalignment increase is not clear, but here we showed that it is not caused by p53 activation or apoptosis. It is possible that Kif20b may help anchor or stabilize midbodies at the apical membrane via cell adhesion junctions until abscission is complete.

Aside from regulating abscission, Kif20b has postmitotic roles in cortical neurons that may contribute to cortical growth. We previously showed that *Kif20b* mutant embryonic cortical neurons in culture display a polarization defect and abnormalities in neurite growth and branching. Thus the deficit in thickness of the cortical plate at birth may be due to reduced neuropil. A key function of Kif20b is to enable tight microtubule packing: Kif20b has microtubule crosslinking activity *in vitro*, and in *Kif20b* mutant neurons, wider neurites and gaps in microtubule bundles were noted (Abaza et al., 2003; McNeely et al., 2017). Therefore, Kif20b may help to crosslink microtubules in both NSC midbodies and neuronal axons.

### Elucidating the heterogeneous mechanisms of brain malformations requires many genetic models and culture systems

Defects in cytokinetic abscission mechanisms could underlie a range of poorly understood microcephalies and related brain malformations. For example, human genetic studies in two different families showed that a very severe prenatal lethal microcephaly/anencephaly is caused by mutations in the gene encoding the key abscission protein Cep55 (OMIM Ref #236500) (Frosk et al., 2017; Bondeson et al., 2017) Cep55 depletion in cell lines causes more severe midbody structural defects and abscission delay than Kif20b depletion does (Zhao et al., 2006). Patients with *Kif20b* loss-of-function mutations have not yet been identified, so it remains to be seen whether the severity of single-cell abscission phenotypes correlates with the severity of brain malformation. Cell lines such as HeLa cells can model some abscission defects, for example midbody maturation, but not others, such as midbody positioning at the apical membrane, p53 activation, or apoptosis. Our studies of the *Kif20b* mouse model provide human geneticists with a candidate gene and cellular markers for syndromes involving peri- or prenatal lethality, microcephaly, craniofacial defects, or microphthalmia. Comparing mouse models for abscission genes with other microcephalic mice as well as human phenotypes will help us understand the heterogeneous etiologies of and potential diagnosis and treatments for these devastating conditions.

## MATERIALS AND METHODS

### Mice

Mouse colonies were maintained in accordance with NIH guidelines and policies approved by the IACUC. Embryos were harvested by caesarean section, and the morning of the vaginal plug was considered embryonic day (E) 0.5. Littermate embryos served as controls for all experiments. The *Kif20b*^*magoo*^ allele, as previously described (Janisch et al., 2013) is maintained on both C57BL/6 and FVB/N backgrounds, and 50/50% mixed background embryos are used for experiments. *Trp53*^*tm1Tyj*^ mice on C57BL/6 background were obtained from The Jackson Laboratory (JAX stock #002101) (Jacks et al., 1994). Mixed BL6/FVB/N background *Bax*^*tm1Sjk*^ mice were a gift from Christopher Deppmann (JAX stock #002994) (Knudson et al., 1995). Sex of embryonic mice were not noted as sex was not a relevant biological variable for these experiments. The specific ages of embryonic mice used is noted in figure legends for each experiment.

### Cortical cell cultures

Cells were dissociated from E12.5 cortices following a protocol adapted from Sally Temple’s lab (Qian et al., 1998). The Worthington Papain Dissociation Kit was used to dissociate cells (Worthington Biochemical Corporation, Cat # LK003150). Cells were cultured in DMEM with Na-Pyruvate, L-Glutamine, B-27, N2, N-acetyl-cysteine and basic Fibroblast Growth Factor (bFGF). After 24 hours, cells were fixed by adding an equal volume of room-temperature 8% PFA for 5 minutes to cell media, followed by removal of media and addition of - 20° cold methanol for 5 minutes.

### Apical slab preparation

Apical slabs were prepared as previously described (Janisch and Dwyer, 2016). The meninges and skull were removed to expose the brain in E13.5 embryos, followed by fixation with 2% PFA for 20 minutes. Next, apical slabs were made by pinching off cortices, flipping so that the apical surface was upright, and trimming to flatten the slab. Slabs were fixed for another 2 minutes with 2% PFA followed by blocking with 5% normal goat serum (NGS) for 1 hr. Primary antibodies were applied for 1 hr. at room temperature and then moved to 4° overnight. The next day, after 3, 10-minute PBS washes secondary antibodies and DAPI were applied at a concentration of 1:200 for 30 minutes. After two more 10 minute PBS washes slabs were coverslipped with VectaShield fluorescent mounting medium (Vector Laboratories Inc., H-1000) and imaged. *z*-stack depth was 8-20 µm and step size was 0.5 µm. Midbodies were considered misaligned if the two halves of the midbody were not in the same *z*-plane or within two adjacent *z*-planes.

### Immunoblotting

Brain lysates were prepared with RIPA lysis buffer (150 mM NaCl, 1% NP40, 0.5% sodium deoxycholate, 0.1% SDS, 50 mM Tris-HCl pH 8) with protease and phosphatase inhibitors. Protein concentration in lysates was determined by bicinchoninic acid (BCA) assay, and 60 µg total protein was loaded per lane on 4-20 gradient% polyacrylamide gels. Proteins were transferred by electroblotting onto a 0.2 µm PVDF membrane overnight at 30 mA. Membranes were blocked in 150 mM NaCl, 100 mM Tris-HCl pH 7.5 and 0.5% Tween 20 (TBST) with 5% dried milk (blocking buffer) for 1 hour. Primary antibodies were incubated with the membrane overnight at 4°C. After three washes, LI-COR IRDye 800 CW goat anti-rabbit IgG and 680 RD goat anti-mouse secondary antibodies were applied (1:10000) in blocking buffer for 1 hour at room temperature. After washing with TBST for 5 minutes, 3 times, immune complexes were visualized using a LI-COR western blot imager.

### Immunostaining

To collect cryosections for IHC, age E14.5 and P0 brains were removed from heads and fixed for 6 and 24 hours, respectively, in 4% PFA, followed by immersion in 30% sucrose in PBS overnight. Next, whole brains were embedded in OTC (Tissue-Tek, 4583) and cryosections were cut at 20 μm thickness and collected on Superfrost Plus slides (Fisher Scientific, 12-550-15). Frozen sections were stored at −80 degrees. Prior to immunostaining, cryosections were warmed to room temperature, then if antigen retrieval was needed, immersed in 10 mM citrate buffer at 95 degrees for 20 minutes. After cooling, sections were blocked in 2% NGS for 1 hour, followed by incubation with primary antibodies overnight at 4° C. The next day, after PBS washes sections were incubated with AlexaFluor secondary antibodies at 1:200 and DAPI at 1:100 for 30 minutes followed by PBS washes and coverslipping with VectaShield fluorescent mounting medium. For IF on coverslips of dissociated cortical progenitors, a similar protocol was used but with primary antibodies applied for 3 hours at room temperature, and antigen retrieval was not used. Coverslips were mounted on Superfrost Plus slides with Fluoro-Gel (Electron Microscopy Sciences, 17985-10).

### Antibodies

Antibodies used in this analysis were rabbit polyclonal anti-human cleaved-caspase 3 (Cell-Signaling 9661S, 1:250), mouse monoclonal anti-rat beta-III-tubulin (Tuj1) (BioLegend 801201, 1:500), rat monoclonal anti-mouse Tbr2 (ebioscience 14-4875, 1:200), rabbit polyclonal anti-mouse Pax6 (BioLegend PRB-278P, 1:200), Aurora B kinase mouse monoclonal anti-rat (BD Biosciences 611082, 1:300), rabbit polyclonal anti-human alpha-tubulin (Thermo Scientific RB-9281-P0, 1:300), rat monoclonal alpha-tubulin (Novus Biologicals NB600-506, 1:300), rabbit monoclonal anti-human PH3 (Cell Signaling 3458, 1:200), chicken polyclonal anti-mouse Nestin (Aves Labs NES, 1:600), rat monoclonal anti-human Ki67 (eBioscience 14-5698, 1:100), mouse monoclonal anti-human p53 (Millipore 05-224, 1:250), rabbit polyclonal anti-mouse p53 (Leica Biosystems NCL-L-p53-CM5p, 1:500), rabbit polyclonal anti-human Beta-Catenin (Sigma-Aldrich SAB4500545 1:1000), and polyclonal rabbit anti Zo-1 (rabbit, Invitrogen 61-7300,1:50). All antibodies were validated for the application used in multiple previous publications.

Images were collected on either a Zeiss Axio ImagerZ1 microscope with AxioCam MRm (Figures 1, 2, 3, 5 6 and 7A-C), a Zeiss AxioObserver fluorescent widefield inverted scope microscope (Figures 1, 2 and 3), an inverted DeltaVision with TrueLight deconvolution microscope and softWoRx Suite 5.5 image acquisition software (Applied Precision) (Figure 7D-J), or a Leica MZ16F microscope with DFC300FX camera (Figure 2, H&E and Figure 4). **Control and mutant fluorescence images were collected with the same exposure times and on the same day.** All image analysis was performed in ImageJ/Fuji and any changes to brightness and contrast were applied uniformly across images. Statistical analyses were performed using Excel (Microsoft) or GraphPad PRISM software. The sample sizes were pre-determined based on our labs previous experience with cortical development analyses and others’ published results. After obtaining pilot data, power analyses were performed if necessary to determine if the number of samples obtained was high enough for the effect size seen. NSC cultures that were unhealthy were not imaged and analyzed, but no other data was excluded from the analysis. No randomization or blinding was used as no experimental manipulation was applied other than genetic knockouts. Genotyping was performed after collection of embryos to determine genetic status. Statistical tests used are specified in each figure legend. For each sample set a statistical test of normality was performed using GraphPad PRISM software. Parametric tests were used when sample sets had a normal distribution and non-parametric tests were used when sample sets did not have a normal distribution. Variance was calculated and was typically similar between groups. N’s listed in figure legends indicate the number of coverslips, brains and litters collected for each experiment. For each brain, at least three sections were imaged, and for each coverslip, at least 5, 20X pictures or 10, 40X pictures were analyzed.

## ACKNOWLEDGEMENTS

We thank Chris Deppmann for use of the *Bax* mutant mouse line. We are grateful to Bettina Winckler, Jing Yu, Xiaowei Lu, Ann Sutherland, Todd Stukenberg and members of their labs, as well as Katrina McNeely for advice and discussion. We are grateful to Madison Hecht, Gabrielle Wolfe, Mackenzie Shannon, Haley Hopkinson and Adriana Ehlers for help with cryosectioning and data analysis. This work was supported by the National Institutes of Health (R01 NS076640 to NDD), the UVA Medical Scientist Training Program (MSTP T32 GM 7267-37) and the UVA Cell and Molecular Biology Training Grant (2T32GM008136-31A1).

---

**Supplemental Figure S1:**
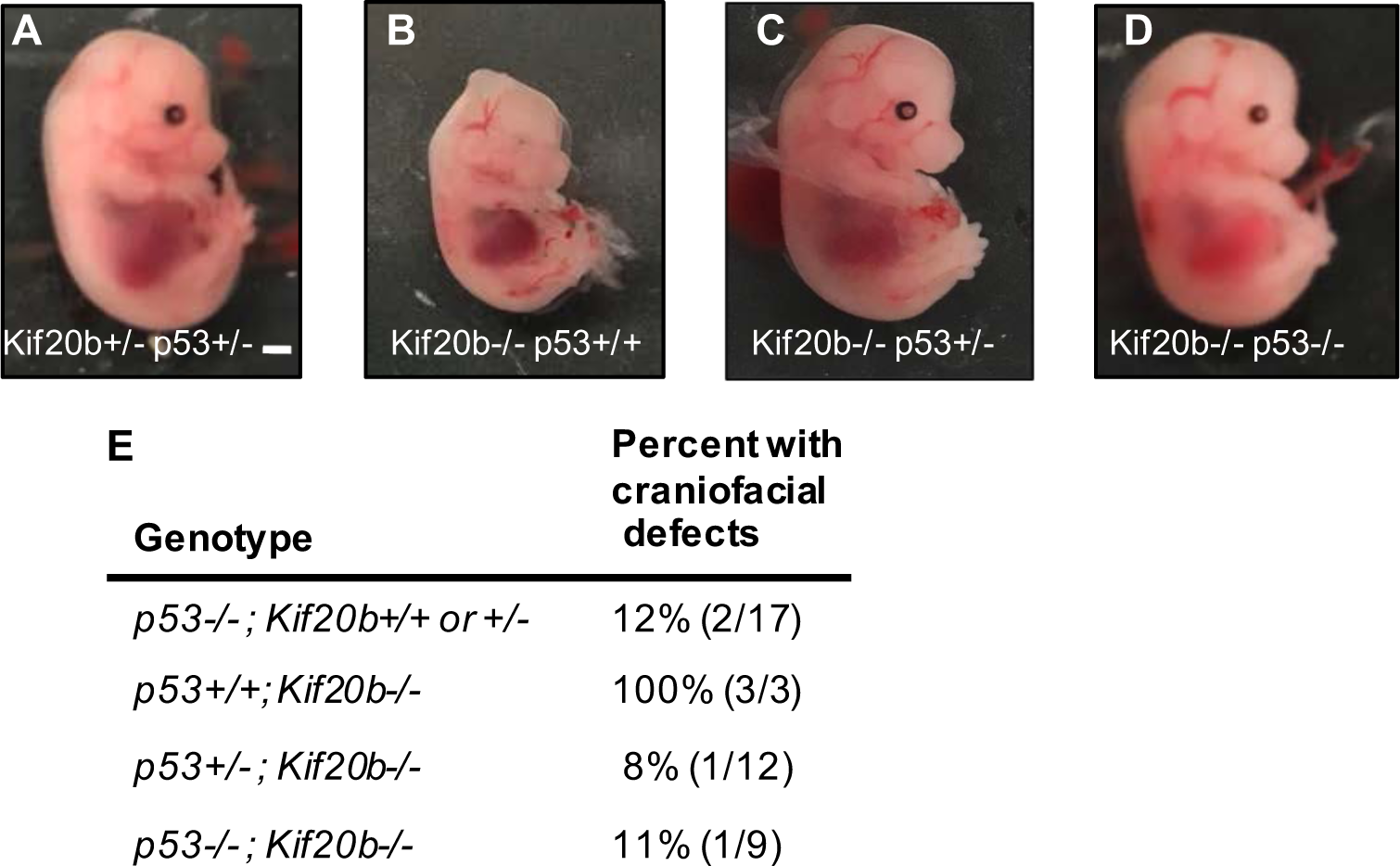
*p53* deletion rescues craniofacial defects in *Kif20b* mutant mice. **A-D.** Representative E14.5 embryos of the indicated genotypes. **E.** E14.5 embryos observed with craniofacial defects including shortened snout, underdeveloped eyes, and/or brain malformations of the indicated genotypes.

**Supplemental Figure S2.**
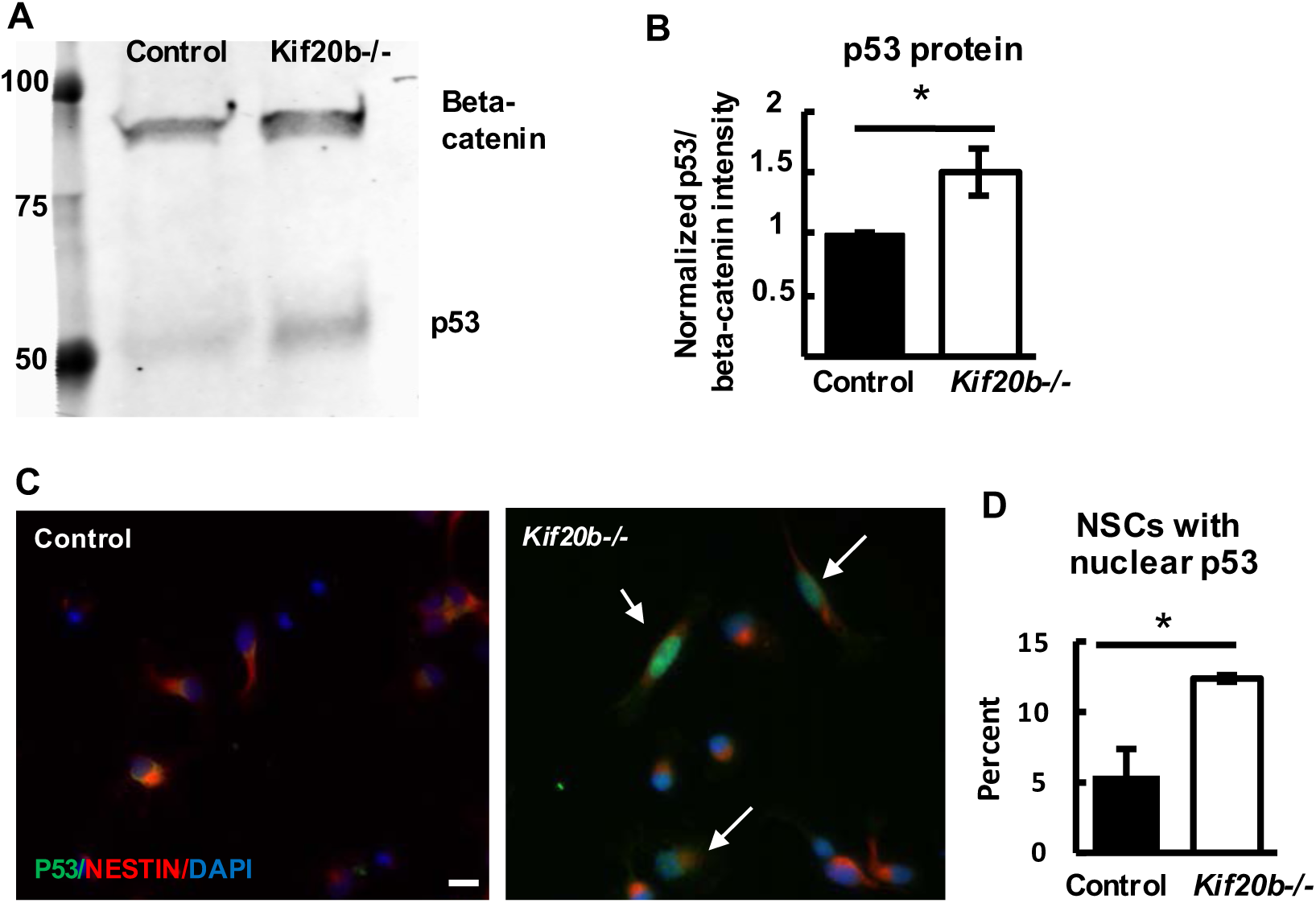
p53 protein level is elevated in *Kif20b-/-* E12.5 brain lysates and dissociated NSCs. **A.** Immunoblot from E12.5 control and *Kif20b-/-* cortical lysates shows bands for p53 (53 kDa) and beta-catenin (95 kDa) as a loading control. **B.** p53 band intensity, normalized to beta-catenin bands, is increased by 50% in *Kif20b-/-* samples. n = 4 blots from 4 embryos each. * p ≤ 0.05, paired ratio t-test. **C.** Representative images of E12.5 dissociated cortical cultures fixed after 24 hrs *in vitro*, and immunostained for p53 (green) and nestin (red) to mark NSCs. Scale bar: 10 μ m. **D.** The average percent of NSCs with detectable nuclear p53 staining is increased more than two-fold in *Kif20b-/-* cultures. n= 3 cultures for each genotype, each from an independent E12.5 brain.

**Supplemental Figure S3.**
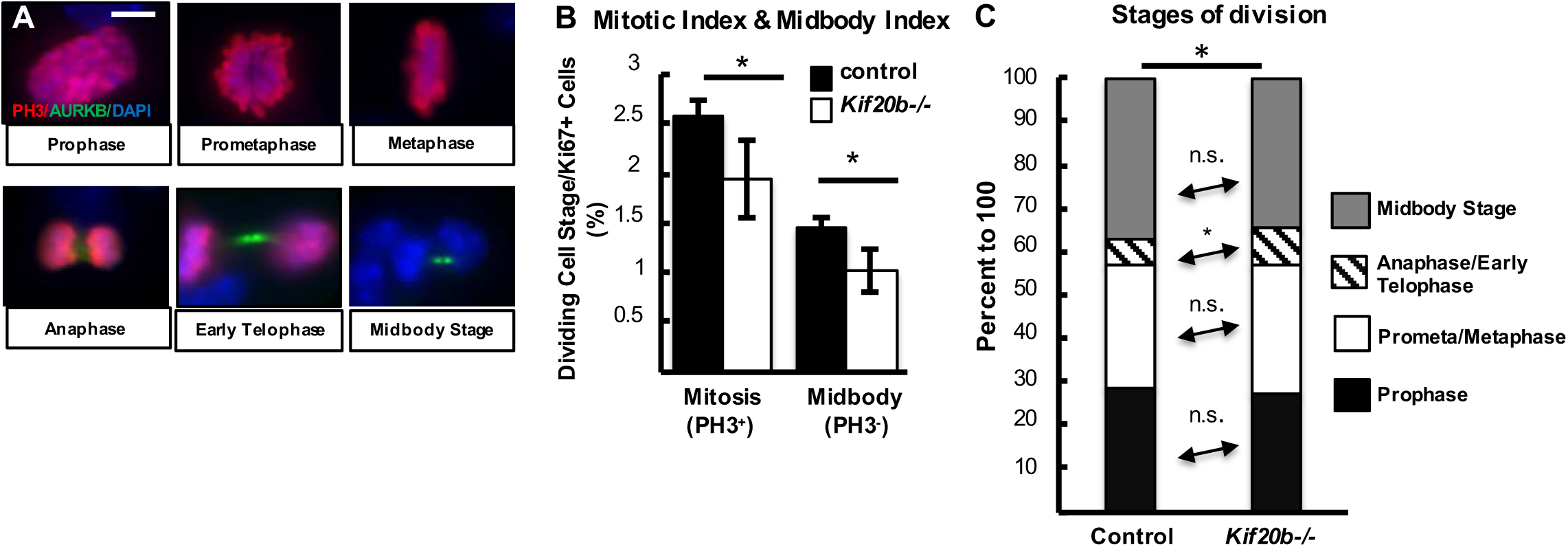
Analyses of mitotic and midbody indices in *Kif20b* mutant NSC cultures. **A.** Examples of E12.5 dissociated cortical NSCs at stages of mitosis and cytokinesis, labeled with PH3 (red) and AurkB (green). **B.** The mean percentages (+/- s.e.m.) of cycling NSCs(Ki67+) in mitosis or midbody stage are slightly reduced in *Kif20b-/-* cultures. **C.** Categorizing the dividing NSCs from B, there is a small increase of *Kif20b-/-* NSCs in anaphase/early telophase. For B and C, n = 3 coverslips from 3 embryos each, with 1499 control and 1421 *Kif20b-/-* Ki67+ cells. * p ≤ 0.05, Student’s t-test for (B), Chi Square and Fisher’s exact test for (C). All cultures dissociated from E12.5 cortices and fixed after 24 hours. Scale bar, 5 μ m.

